# The iPSYCH2012 case-cohort sample: New directions for unravelling genetic and environmental architectures of severe mental disorders

**DOI:** 10.1101/146670

**Authors:** Carsten Bøcker Pedersen, Jonas Bybjerg-Grauholm, Marianne Giørtz Pedersen, Jakob Grove, Esben Agerbo, Marie Bækvad-Hansen, Jesper Buchhave Poulsen, Christine Sæholm Hansel, John J. McGrath, Thomas Damm Als, Jacqueline I. Goldstein, Ben M Neale, Mark J Daly, David M. Hougaard, Ole Mors, Merete Nordentoft, Anders D. Børglum, Thomas Werge, Preben Bo Mortensen

**Affiliations:** iPSYCH, The Lundbeck Foundation Initiative for Integrative Psychiatric Research, Denmark; National Centre for Register-Based Research, Aarhus University, Business and Social Sciences, Aarhus, Fuglesangs Alle 4, 8210 Aarhus V, Denmark; Centre for Integrated Register-based Research, CIRRAU, Aarhus University, Aarhus, Denmark; Department for Congenital Disorders, Statens Serum Institut, Copenhagen, Denmark; Department of Biomedicine and iSEQ, Centre for Integrative Sequencing, Aarhus University, Denmark; BiRC-Bioinformatics Research Centre, Aarhus University, Aarhus, Denmark; Queensland Brain Institute, The University of Queensland, St Lucia, Australia; Queensland Centre for Mental Health Research, The Park Centre for Mental Health, Wacol, Australia; Analytic and Translational Genetics Unit (ATGU), Department of Medicine, Massachusetts General Hospital and Harvard Medical School, Boston, Massachusetts, USA; Program in Medical and Population Genetics, Broad Institute of Harvard and MIT, Cambridge, Massachusetts, USA; Stanley Center for Psychiatric Research, Broad Institute of Harvard and MIT, Cambridge, Massachusetts, USA; Psychosis Research Unit, Aarhus University Hospital, Risskov, Denmark; Copenhagen University Hospital, Mental Health Centre Copenhagen, Capital Region of Denmark, Denmark; Institute of Biological Psychiatry, Mental Health Centre Sct. Hans, Capital Region of Denmark, Copenhagen University Hospital, Denmark

## Abstract

The iPSYCH consortium has established a large Danish population-based Case-Cohort sample (iPSYCH2012) aimed at unravelling the genetic and environmental architecture of severe mental disorders. The iPSYCH2012 sample is nested within the entire Danish population born 1981-2005 including 1,472,762 persons. This paper introduces the iPSYCH2012 sample and outlines key future research directions. Cases were identified as persons with schizophrenia (N=3,540), autism (N=16,146), ADHD (N=18,726), and affective disorder (N=26,380), of which 1928 had bipolar affective disorder. Controls were randomly sampled individuals (N=30,000). Within the sample of 86,189 individuals, a total of 57,377 individuals had at least one major mental disorder. DNA was extracted from the neonatal dried blood spot samples obtained from the Danish Neonatal Screening Biobank and genotyped using the Illumina PsychChip. Genotyping was successful for 90% of the sample. The assessments of exome sequencing, methylation profiling, metabolome profiling, vitamin-D, inflammatory and neurotrophic factors are in progress. For each individual, the iPSYCH2012 sample also includes longitudinal information on health, prescribed medicine, social and socioeconomic information and analogous information among relatives. To the best of our knowledge, the iPSYCH2012 sample is the largest and most comprehensive data source for the combined study of genetic and environmental aetiologies of severe mental disorders.

## Introduction

The fundamental nature of mental disorders remains poorly understood. Twin studies have suggested heritability estimates around 80% for schizophrenia, bipolar affective disorder, autism and ADHD, with slightly lower heritability estimates for depression^1^. Considerable progress in psychiatric genetics has been made in recent years, based on large samples and international collaborations, e.g., through the pivotal efforts of the Psychiatric Genomics Consortium (PGC)^2^. We can expect that larger samples will reveal new insights to common and rare variants underpinning mental disorders^3^.

Several environmental factors have been associated with mental disorders. Many environmental factors influencing pre - and postnatal development are associated with schizophrenia, bipolar affective disorder, autism, and ADHD, and furthermore, adverse life circumstances increase the risk of mental disorders. Gene - environment synergism contributes to the aetiology of these disorders, but suitable datasets to explore this important field of research have been lacking. To understand the impact of genes and environments over the life course, large and truly population-based longitudinal cohort studies are required^4^.

As part of the Lundbeck Foundation Initiative for Integrative Psychiatric Research (iPSYCH: http://iPSYCH.au.dk/) a large case-cohort study has commenced. In most countries, it would not be logistically feasible to compile large, representative population-based samples. In Denmark, the existence of (a) a universal public health care system free of charge, (b) several national longitudinal registers, and (c) strict ethical and data protection legislation required to safeguard the privacy of study participants, has provided a remarkable research platform^5^. Recent technological developments and a new legal framework for use of bio-banked material for research have created similar possibilities for genetic research.

The vision of iPSYCH was to leverage these combined resources, considering the entire national cohort as our study population. We utilized information on individuals with a diagnosis of selected mental disorders (N=57 377) and a randomly sampled cohort^6,7^ of the general population (N=30 000). The sample is known as the iPSYCH Danish case-cohort study, in short iPSYCH2012. We used neonatal dried blood spots from the Danish Neonatal Screening Biobank to investigate detailed genetic and biomarker information, some of which are markers of environmental exposures. The rich Danish population-based registers were used to add information on all individuals and all their relatives. Thus, we created a comprehensive data source for the combined study of genetic and environmental aetiologies of severe mental disorders. Within the iPSYCH2012 sample, currently around 77 500 individuals have been array genotyped and around 20 000 have been whole exome sequenced. Ten thousand samples have been analysed for ranges of cytokines and neurotrophic factors. Epigenetic and metabolome data from several thousand samples are emerging. For the entire sample and their relatives, detailed longitudinal information related to health, prescribed medicine, social and socioeconomic information exists. This paper provides a general overview of the sample design, and outlines future research.

### The overall design

Individuals diagnosed with schizophrenia, mood disorders, bipolar affective disorder, autism and ADHD were identified through linkage between Danish population-based registers along with a random sample of the same population that supplied the cases^8^. Dried blood spots for virtually all individuals were retrieved from the Danish Neonatal Screening Biobank and processed for genotyping. The design includes the ability to efficiently analyse prospectively collected cohort data within the iPSYCH case-cohort sample^8^. This particular design provides several advantages: Because the cohort is randomly selected from the entire population, we are able to generate unbiased absolute risks and incidence rates, and to estimate the effect sizes of genetic markers on risk of mental disorders which is representative of the entire Danish population. To date, most genetic and epidemiological studies are based on convenient case-control samples, which are prone to biases^8,9^. The iPSYCH2012 sample was preceded by four smaller Danish samples^10–17^, all aiming to investigate the potential interplay between genes and the environment. Collectively, these forerunners informed on the best possible study design to use in the iPSYCH2012 sample (Supplementary text 1). The following three paragraphs describe the resources and methods used to identify individuals included in the iPSYCH2012 sample.

### Selecting the study base

The Danish Civil Registration System was established in 1968^18^, where all people alive and living in Denmark were registered. It includes information on the unique personal identification number, sex, date and place of birth, parents' identifiers, and continuously updated information on emigration and death. The personal identification number is used in all national registers enabling accurate linkage within and between registers. The study base included all singleton births with known mothers born between 1^st^ of May 1981 and 31^st^ of December 2005 who were alive and resided in Denmark at their 1^st^ birthday (N= 1 472 762 persons). Selecting births in this period ensures individual samples to be retrieved in the Danish Neonatal Screening Biobank and reasonable distribution of cases and cohort members for all birth years. All residents are registered in the Danish Civil Registration System irrespective of health, income, receipt of social benefits, employment and other socioeconomic characteristics^18^.

### Diagnoses of mental disorders

Persons within the study base were linked via their personal identifier to the Danish Psychiatric Central Research Register^19^ to obtain information on mental disorders. The Danish Psychiatric Central Research Register was computerized in 1969 and contains data on all admissions to Danish psychiatric in-patient facilities. Information on outpatient visits was included from 1995 onwards. From 1994 onwards, the International Classification of Diseases, 10th revision, Diagnostic Criteria for Research (ICD-10-DCR) was used for diagnostic classification^20^. All persons within the study base who had a diagnosis of schizophrenia, bipolar disorder, affective disorder, autism and ADHD were included (Table 1). All patients were identified in the Danish Psychiatric Central Research Register. At the time of linkage, the register contained all psychiatric contacts until 31^st^ December 2012. Table 1 summarizes the number of individuals across the diagnostic groups.

**Table 1:** Number of persons included in iPSYCHs population-based sample of the Danish population born 1981-2005

| Group membership<sup>a</sup> | ICD10 diagnoses<sup>b</sup> | Follow-up period | Number of persons |  |  |
| --- | --- | --- | --- | --- | --- |
|  |  |  | Initial Sample | Blood spots available in Biobank and sent to Broad | Passed Sample QC |
| Schizophrenia<sup>c</sup> | F20 | 2009-2012 | 3540 | 2830 | 2738 |
| Bipolar disorder | F30-F31 | 1994-2012 | 1928 | 1488 | 1452 |
| Affective disorder | F30-F39 | 1994-2012 | 26380 | 23532 | 22809 |
| Autism | F84.0, F84.1, F84.5, F84.8 or F84.9 | 1994-2012 | 16146 | 15418 | 14812 |
| ADHD | F90.0 | 1994-2012 | 18726 | 17835 | 17249 |
| Any case | All ICD codes listed above |  | 57377 | 52867 | 51101 |
| Population-based cohort | Random sampling, i.e. disregarding any diagnostic information |  | 30000 | 28650 | 27605 |
| Total number of persons |  |  | 86189 | 80422 | 77639 |

<sup>a</sup> Groups are not mutually exclusive.
<sup>b</sup> Initial ICD10 diagnosis used to select case groups. Identification was performed through linkage to the Danish Psychiatric Central Research Register. At the time of linkage, the register contained all contacts up until 31/12/2012.
<sup>c</sup> Include all persons first diagnosed with schizophrenia during the period from 1/1/2009 to 31/12/2012 and persons first diagnosed with schizophrenia before 1/1/2009 that were not already genotyped (N=923). A total of 1887 persons first diagnosed with schizophrenia before 1/1/2009 were previously genotyped (see supplementary text 1).

### Selecting the population-based cohort

Among the 1 472 762 persons included in the study base, a total of 30 000 persons were chosen uniformly at random (Table 1) corresponding to 2.04% of the study base (=30 000/1 472 762). Because the cohort members were chosen randomly, some cohort members may also have the disorders of interest^6,7^. Thus, the cohort selected is representative of the entire Danish population born in the same period^18^. In addition, the cohort members are at risk of developing the disorder of interest during follow-up, while controls are typically conditioned to be healthy until the study ends^21^. We have thereby identified the individuals to be included in the iPSYCH2012 sample. Next we describe the enrichment with genetic and other biomarker data.

### The Danish Neonatal Screening Biobank

Blood spots for individuals included in the iPSYCH2012 sample were retrieved from the Danish Neonatal Screening Biobank within the Danish National Biobank^22^. This facility stores dried blood spot samples taken from practically all neonates born in Denmark since 1 May 1981 and stored at −20 degree Celsius. These samples were collected primarily for diagnosis of congenital disorders. The samples are stored for follow-up diagnostics, screening, quality control and research. At time of blood sampling (4-7 days after birth), parents are informed in writing about the neonatal screening and that the blood spots are stored in the Danish Neonatal Screening Biobank and can be used for research, pending approval from relevant authorities. The parents are also informed about how to prevent or withdraw the sample from inclusion in research studies. Genotyping was based on two blood spot punches of 3.2mm, equivalent to 6 μ! of whole blood^22^. Neonatal dried blood spot samples may be challenging to analyse due to the very limited amount of biological material available, the nature of dried whole blood on filter paper and decades of storage. Special assays may be required and multi-analyte measurements are preferred to get as much information of the samples as possible. The neonatal dried blood spots is also suitable for next generation sequencing^23^, DNA methylation profiling^24^, metabolome profiling, vitamin D^25^, multiplex measurements of cytokines ^26^, antibodies to infectious agents^12^, and whole transcriptome analysis through microarray and RNA-seq^27^. Importantly, these measurements are made in samples drawn few days after birth, meaning that case-control differences cannot be ascribed to disease-related confounders as medication, alcohol or substance use, smoking or the disease state itself.

*Preparation of samples for genotyping and sequencing from the Danish neonatal screening biobank* DNA was extracted and whole genome amplified at the Statens Serum Institut following previously established procedures^28,29^. The sample flow is described in Figure 1.

**Figure 1:**
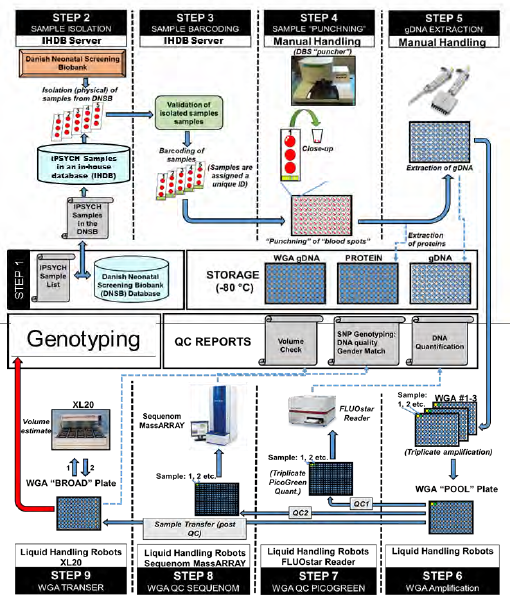
The selected samples were correlated with their DNSB identifiers and entered into an in-house developed selection database (Step 1 and 2). Sample identities were then validated and assigned a pseudonymized unique ID (Step 3) before cutting two discs of 3.2mm of dried blood into a 96 well PCR plate (Step 4). Proteins were washed of the blood spots and stored at −80° C before DNA was extracted using Extract-N-Amp Blood PCR Kit (Sigma-Aldrich, St. Louis, United States of America) (Step 5). DNA was amplified in amplified in triplicates using REPLI-g (Qiagen, Hilten, Germany) and combined to a single sample (Step 6). Finally concentrations were quantified using Quant-iT picogreen (Invitrogen, California, United States of America) (Step 7) and a genetic fingerprint established using the iPLEX pro Sample ID panel (Agena Bioscience, Hamburg, Germany) (Step 8) before aliquoting a fraction of the sample for genotyping (Step 9).

### Array Genotyping and Quality Control

Samples were processed at the Broad Institute (Boston, Massachusetts, United States of America) using the Infinium PsychChip v1.0 array (Illumina, San Diego, California, United States of America) in accordance with the manufacturer's instructions^30^. Genotyping was conducted in 25 waves. Variant calls were trained using GenTrain2 (Illumina) on the first wave (4 146 samples) using the PsychChip 15048346 B manifest and GenomeStudio version v2011.1. Following autoclustering, loci were manually curated if they had a call frequency below 90%, GenTrain scores below 0.5 or cluster separation below 0.2. During this processing, 3 890 loci were excluded and 928 were manually modified. The resulting GenTrain was used to produce GenCall variant calls used for sample level quality control of the entire cohort^31^. Samples with call rates below 95% (N=2 270) were designated to fail sample QC. Sex was inferred using heterozygosity on chromosome X; below 20% in males; above 20% in females. Sex obtained from genotyping was compared to the sex recorded in the Danish Civil Registration System and mismatches were excluded. It is extremely unlikely to observe errors in recorded sex in the Danish Civil Registration System^18^. About 0.25% (N=224) of the sample did not match the expected sex. Half of the failures (N=119) were due to abnormal structural variation on chromosome X (aneuploidy and loss of heterozygosity). The other half were due to sample mix-ups (N=103). In this paper we describe the sample QC only and not the subsequent SNP QC that is likely to change between studies.

### Probe remapping

All probe sequences were queried against an HG19 database using a nucleotide version of the Basic Local Alignment Search Tool (BLAST). The Blast results were compared to the original array manifest, an Illumina update to the array manifest, and the Broad Institute updates to the manifest. The genomic coordinates matched between the BLAST results and the existing manifests for 95.12% of probes. 2.23% of probes were updated based on the new BLAST results. 2.11% retained their original mapping. The remaining 0.54% were split between the Broad Institute reference and the Illumina update, or the probe was removed from the dataset (Supplementary Table 1).

### Improving variant calls

GenCall^31^, Birdseed^32^ and zCall^33^ were used supplementary to improve variant calls. GenCall and Birdseed are genotype calling algorithms best suited for common variants, while zCall is a post-processing step for GenCall to improve genotype calling for rare variants. Approximately half of the probes on the array are common variants (MAF ≥ 0.05) while the other half are rare variants (MAF < 0.05). A large subgroup of the rare variants are non-polymorphic within the cohort. A consensus genotype call was made from the three calling algorithms described in Supplementary text 2 using PLINK^34,35^.

### Ethical framework

The Danish Scientific Ethics Committee, the Danish Health Data Authority, the Danish data protection agency and the Danish Neonatal Screening Biobank steering committee, approved this study. This is in keeping with the strict ethical framework and the Danish legislation protecting the use of these samples^22,36^. Permission has been granted to study genetic and environmental factors for the development and prognosis of mental disorders. To unravel the foundation of severe mental disorders, it is central that this rich data source is accessible to the international research community to the largest extent possible. It is paramount to protect the privacy of the individuals included in the study. Due to the sensitive nature of these data, individual level data can be accessed only through secure servers where download of individual level information is prohibited^37^. Current possibilities for access to the iPSYCH2012 sample include access through Genome Denmark (http://www.genomedenmark.dk/), ComputerOme (https://computerome.dtu.dk/) and Statistics Denmark (http://www.dst.dk/en/TilSalg/Forskningsservice).

### Baseline characteristics

Table 2 shows baseline characteristics of the 86 189 individuals included in the iPSYCH2012 sample. Among these individuals, 77 639 (90%) passed sample QC. In the cohort group, males constituted 51% in both the initial and in the QC'ed sample. The following numbers refer to the initial sample: Overall, 26 380 individuals were included due to suffering from an affective disorder. Among individuals with affective disorder, 543 individuals were incidentally also among the cohort members, i.e. the 2.03% random sample of the study base. Overall, 28 812 (96.04%) of the 30 000 cohort members had none of the five psychiatric diagnoses until 2012. A total of 49 737 (86.68%) cases and 25 159 (83.86%) cohort members were native Danes. The largest second-generation immigrant group was persons having one or both parents born in Europe followed by one or both parents born in Scandinavia.

Comparing the percentage of cases included in the initial sample to the percentage of cases passing the QC revealed no systematic deviations across selected baseline characteristics (Table 2). Comparing the percentage of cohort members included in the initial sample to the percentage of cohort members passing QC also revealed no systematic deviations across baseline characteristics.

**Table 2:** Baseline characteristics of the iPSYCH2012 case-cohort

|  | Initial sample, N=86189 |  | Passed sample QC, N=77639 |  |
| --- | --- | --- | --- | --- |
|  | Cases<sup>a</sup>, N=57377 | Cohort<sup>a</sup>, N=30000 | Cases<sup>a</sup>, N=51101 | Cohort<sup>a</sup>, N=27605 |
| Sex |  |  |  |  |
| Male | 32243 (56.19%) | 15308 (51.03%) | 29048 (56.84%) | 14090 (51.04%) |
| Female | 25134 (43.81%) | 14692 (48.97%) | 22053 (43.16%) | 13515 (48.96%) |
| Year of birth |  |  |  |  |
| 1981-1985 | 11909 (20.76%) | 4858 (16.19%) | 9409 (18.41%) | 4175 (15.12%) |
| 1986-1990 | 14506 (25.28%) | 5838 (19.46%) | 12766 (24.98%) | 5361 (19.42%) |
| 1991-1995 | 14521 (25.29%) | 6597 (21.99%) | 13508 (26.43%) | 6190 (22.42%) |
| 1996-2000 | 10306 (17.96%) | 6461 (21.54%) | 9796 (19.17%) | 6182 (22.39%) |
| 2001-2005 | 6144 (10.71%) | 6246 (20.82%) | 5622 (11.00%) | 5697 (20.64%) |
| Diagnosis<sup>b</sup> |  |  |  |  |
| Schizophrenia | 3540 (6.17%) | 90 (0.30%) | 2738 (5.36%) | 65 (0.24%) |
| Bipolar affective disorder | 1928 (3.36%) | 42 (0.14%) | 1452 (2.84%) | 31 (0.11%) |
| Affective disorder | 26380 (45.98%) | 543 (1.81%) | 22809 (44.64%) | 467 (1.69%) |
| Autism | 16146 (28.14%) | 324 (1.08%) | 14812 (28.99%) | 309 (1.12%) |
| ADHD | 18726 (32.64%) | 387 (1.29%) | 17249 (33.75%) | 360 (1.30%) |
| None of the above | 0 (0.00%) | 28812 (96.04%) | 0 (0.00%) | 26538 (96.13%) |
| Parental origin<sup>c</sup> |  |  |  |  |
| Denmark | 49737 (86.68%) | 25159 (83.86%) | 44284 (86.66%) | 23213 (84.09%) |
| Africa | 619 (1.08%) | 446 (1.49%) | 566 (1.11%) | 400 (1.45%) |
| America | 472 (0.82%) | 228 (0.76%) | 419 (0.82%) | 204 (0.74%) |
| Asia | 731 (1.27%) | 672 (2.24%) | 648 (1.27%) | 616 (2.23%) |
| Australia | 51 (0.09%) | 29 (0.10%) | 47 (0.09%) | 29 (0.11%) |
| Europe excl Scandinavia | 2508 (4.37%) | 1557 (5.19%) | 2228 (4.36%) | 1406 (5.09%) |
| Greenland | 341 (0.59%) | 162 (0.54%) | 306 (0.60%) | 148 (0.54%) |
| Middle East | 666 (1.16%) | 625 (2.08%) | 604 (1.18%) | 568 (2.06%) |
| Scandinavia excl Denmark | 1168 (2.04%) | 600 (2.00%) | 1047 (2.05%) | 549 (1.99%) |
| Mixed parentage | 221 (0.39%) | 166 (0.55%) | 190 (0.37%) | 146 (0.53%) |
| Unknown | 863 (1.50%) | 356 (1.19%) | 762 (1.49%) | 326 (1.18%) |
| Birth weight |  |  |  |  |
| <1500 g | 475 (0.83%) | 145 (0.48%) | 381 (0.75%) | 125 (0.45%) |
| 1500-2499 g | 2613 (4.55%) | 924 (3.08%) | 2197 (4.30%) | 808 (2.93%) |
| 2500-3499 g | 26735 (46.60%) | 13302 (44.34%) | 23758 (46.49%) | 12245 (44.36%) |
| 3500-4499 g | 25500 (44.44%) | 14491 (48.30%) | 22918 (44.85%) | 13425 (48.63%) |
| ≥4500 g | 1739 (3.03%) | 954 (3.18%) | 1600 (3.13%) | 883 (3.20%) |
| Missing information | 315 (0.55%) | 184 (0.61%) | 247 (0.48%) | 119 (0.43%) |
| Gestational age |  |  |  |  |
| <35 weeks | 1342 (2.34%) | 458 (1.53%) | 1105 (2.16%) | 397 (1.44%) |
| 35-36 weeks | 1907 (3.32%) | 857 (2.86%) | 1648 (3.22%) | 776 (2.81%) |
| 37-38 weeks | 8650 (15.08%) | 4305 (14.35%) | 7754 (15.17%) | 3998 (14.48%) |
| 39-40 weeks | 29768 (51.88%) | 15835 (52.78%) | 26642 (52.14%) | 14640 (53.03%) |
| ≥41 weeks | 14995 (26.13%) | 8239 (27.46%) | 13440 (26.30%) | 7594 (27.51%) |
| Missing information | 715 (1.25%) | 306 (1.02%) | 512 (1.00%) | 200 (0.72%) |
| Apgar score (5 minutes) |  |  |  |  |
| 1-5 | 238 (0.41%) | 113 (0.38%) | 204 (0.40%) | 107 (0.39%) |
| 6-7 | 617 (1.08%) | 224 (0.75%) | 523 (1.02%) | 199 (0.72%) |
| 8-9 | 3460 (6.03%) | 1712 (5.71%) | 3083 (6.03%) | 1573 (5.70%) |
| 10 | 52284 (91.12%) | 27590 (91.97%) | 46666 (91.32%) | 25445 (92.18%) |
| Missing information | 778 (1.36%) | 361 (1.20%) | 625 (1.22%) | 281 (1.02%) |

<sup>a</sup> Persons who were both selected as cases and cohortes constitute 1188 persons in the initial sample and 1067 persons in the QC'ed sample. These individuals were counted both in the case and cohort group.
<sup>b</sup> Psychiatric diagnosis at time of selection as described in the method section. Some persons may both be selected in the case and cohort group, have more than one diagnoses, and may later have additional diagnoses.
<sup>c</sup> Persons with both parents born in Denmark were classified as having Danish parental origin. Persons having a Danish-born parent and a foreign born parent were classified as the foreign-born parents region of birth. Persons having two foreign-born parents born in the same region were classified with that region. Persons having two foreign-born parents born in different regions were classified as mixed.

### Visualization of genetic data by foreign parental origin

To visualize population substructure for the genetic data, a principle component analysis was conducted. Pair-wise relatedness coefficients and identical by descent probabilities were estimated using plink v1.90b3v^34,35^ after removal of INDELs and SNPs within long-range disequilibrium regions^38^ or with a missing-rate above 0.001. Relatedness categories were assigned to all pairwise combinations of individuals based on their probabilities of sharing 0, 1 or 2 alleles identical by descent as implemented in GWASTools R-package^39^. One individual from each pair of individuals with a relatedness coefficient above 0.09 was excluded at random. Eigenvectors were inferred using EIGENSOFT version 6.1.4^40^ on the relatedness-pruned set of individuals, and a subset of SNPs pruned for Linkage Disequilibrium (*r^2^* <= 0.05) and filtered for Minor Allele Frequency (MAF >= 0.01). All individuals were subsequently projected onto eigenvectors inferred using the relatedness-pruned set of individuals. Second, using information on parental country of birth recorded in the Danish Civil Registration System, and independently of the genetic data, individuals were classified as both parents born in Denmark or parental foreign country of birth (one or both parents born abroad). Parental foreign country of birth was subdivided according to the geographical region of birth of the foreign parent, i.e. Africa, America, Asia, Australia, Europe (excl. Scandinavia), Greenland, Middle East and Scandinavia^15,41^. The principal component scatterplot shows a clear correspondence between the first two principal components based on the genome-wide SNP genotypic data and parental country of birth as registered in the Danish Civil Registration System (Figure 2). Note, only individuals born in Denmark were included in the sample. Individuals born in Denmark to parents born in other Scandinavian countries clustered together with Danish-born individuals with Danish-born parents, as expected. Within each foreign parental region of birth, individuals with two foreign parents were as anticipated more genetically divergent compared to those having only one foreign-born parent. This finding provides strong evidence of internal validity for the processed genotypes.

**Figure 2:**
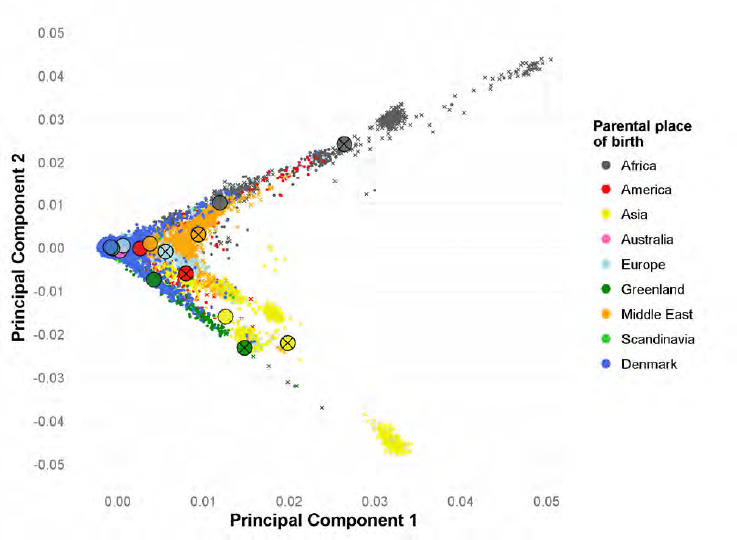
Scatterplot of the first two principal components coloured according to parental region of birth. Big circles indicate mean values for the given parental group. Crosses indicates both parent born abroad within the region indicated by the colour. Absence of cross indicate one Danish born parent and one parent born in the region indicated by the colour. Persons with unknown information on parental region of birth (N=1088) and mixed parentage are not shown (N=366).

### Perspectives

The large iPSYCH2012 sample will provide a solid foundation for a range of studies in decades ahead. We have completed genotyping, and plans are advanced for a range of other analyses, including an update and major expansion with cases diagnosed since 2012 as well as including new diagnostic case groups. The sample is thus not only a rich database for research in the current version - it also constitutes a logistic and organizational framework for future studies, although each new study will require relevant ethical permissions. Most other genetic studies are based on samples of convenience rather than utilizing true population-based samples. To our knowledge, no large-scale population-based sample with GWAS data exists elsewhere. In particular, we are confident that the iPSYCH2012 sample provides an important resource to explore novel ways to combine genetic, phenotypic, and environmental factors. Phenotypic and environmental factors are readily available through record linkage between the numerous Danish registers, or assayed from neonatal dried blood spots.

Access to high quality, population-based person-linked registers, has enabled major contributions to psychiatric epidemiology. For example, researchers have documented key risk factors within psychiatric epidemiology, e.g. urban birth^42^, paternal age^43^, psychiatric family history^44^, life-time risk^45^, infections^10,11,12,13,14^, neonatal vitamin D deficiency^46^, socio-economic adversity^47^, treatment resistant schizophrenia^48^, pharmacological treatment^49^, suicide^50^ and excess mortality^51^. Key features such as the avoidance of selection bias and control of multiple confounders have been important aspects of these studies. However, genetic studies have traditionally not had access to population-based samples, with cases often recruited from multiplex families, or convenient samples of prevalent cases in contact with mental health services. The iPSYCH2012 sample includes a large representative sample of severe mental disorders from a representative sampling frame. The possibility to link the iPSYCH2012 sample to the comprehensive and high quality Danish population-based registers offers researchers unique possibilities to study the interplay between the genetic factors, and variables from the environment, and variables related to health^19,52–54^, mortality, income and social and socioeconomic characteristics^55–57^. Genetic association studies are by default observational studies, subject to many of the same sources of bias and confounding as other epidemiological studies^9^. Therefore, we believe our samples can assist the assessment of the potential impact of such biases and especially lack thereof, and point towards new avenues of research. For example, it has been shown that the genetic associations with schizophrenia identified in the seminal PGC paper^58^ were stronger in more chronic cases than in first episode cases^59^. This may suggest that, in future studies, the genetic architecture of schizophrenia could perhaps be refined to identify genes particularly associated with the risk of developing disease, and genes particularly predicting a chronic course, something that could have important preventive and clinical implications. Such future studies will benefit from the continued dialogue between epidemiological studies as iPSYCH and the large-scale studies available only through collaboration in international consortia.

The iPSYCH2012 sample will be able to leverage SNP-derived, genome-wide metrics such as disease-specific polygenic risk scores^60–62^. These provide a continuous measure of liability (rather than a categorical measure of family history), which will greatly enhance our ability to combine genetic, environmental and phenotypic data in disease prediction. We have found higher polygenic loading for schizophrenia in both cases and controls with family histories of mental disorders^63^. Also 48% of the effect associated with family history of psychoses was mediated through the polygenic risk score for schizophrenia^64^. To further explore the association between the risk of schizophrenia and the polygenetic risk score for schizophrenia, we have investigated the interplay with infections^65^, treatment resistant schizophrenia^66^, chronicity of schizophrenia^59^, and mortality and suicidal behaviour^67^.

Since the initiation of the iPSYCH2012 sample other related Danish projects have built on the same framework as that used within iPSYCH, e.g., anorexia (5 703 cases), obsessive-compulsive disorder (7 747 cases), conduct disorder (4 205 cases) and hyperkinetic conduct disorder (3 690 cases). Also twin samples are in progress within iPSYCH. All samples gain power in utilizing cohort members within the iPSYCH2012 sample while also contributing to the unique possibilities of the iPSYCH2012 sample.

### Strengths and limitations

Identification of cases within the iPSYCH2012 sample is based on contacts to in - and out-patient psychiatric departments and visits to psychiatric emergency care units in a nation where treatment is provided through the government healthcare system free of charge, and where no private psychiatric hospitals exist. Financial factors are thus less likely to influence pathways to healthcare in Denmark compared to many other nations^68^. Unlike samples of convenience, the iPSYCH2012 sample is representative of the Danish population irrespectively of (a) recall bias, (b) emigration or death before sampling, (c) institutional care, (d) imprisonment, (e) being homeless, (f) health, and (g) socioeconomic status^18^. In contrast to most genetic studies, the iPSYCH2012 sample also provides the unique possibility to explore the potential impact of the longitudinal trajectory on causes and outcomes of mental disorders.

While register-based studies like the current study cannot identify persons with untreated disorders or disorders treated in primary health care only, the major strength of the iPSYCH2012 sample approach is the comprehensive clinical assessment of all mental disorders treated in secondary healthcare in a nationwide population. Validation of the Clinician-derived key diagnoses (schizophrenia, single depressive episode, affective disorder, ADHD, autism) has been carried out with good results^69–74^. However, it is a limitation that we are not allowed to re-contact individuals for any reason.

## Conclusion

The iPSYCH2012 case-cohort sample offers a unique foundation for psychiatric research in future decades. It leverages the strengths of several comprehensive Danish health registers. It will accelerate psychiatric research in preventing and treating severe mental disorders. In a Science article titled *'The epidemiologist's dream: Denmark'* ^75^, the ability to examine genomic material in Danish neonatal blood spots was described as a 'gold mine'. iPSYCH are now transforming the 'epidemiologist's dream' into reality.

## Acknowledgements

### Funding

This study was supported by The Lundbeck Foundation (grant no R102-A9118 and R155-2014-1724), Denmark; the Stanley Medical Research Institute; an Advanced Grant from the European Research Council (project no: 294838); the Stanley Center for Psychiatric Research at Broad Institute and Centre for Integrated Register-based Research at Aarhus University. This research has been conducted using the Danish National Biobank resource, supported by the Novo Nordisk Foundation. Professor John J. McGrath is supported by grant APP1056929 from the John Cade Fellowship from the National Health and Medical Research Council and the Danish National Research Foundation (Niels Bohr Professorship).

We thank Betina Trabjerg, National Centre for Register-Based Research, Aarhus University, School of Business and Social Sciences, Aarhus, Denmark for technical help in producing the principal component plot.

We are indebted to the late Mads Vilhelm Hollegaard for his contribution to make sample material accessible for analysis from the Danish Neonatal Screening Biobank. Mads' pioneering work will be used in this and future studies.

### Conflicts of interests

The funders were not involved in this study nor the decision to publish this paper. All other authors report no conflicts of interests.

Supplementary information is available at MP's website.

## Supplementary text 1

The ÌPSYCH2012 had four precursors; all aiming at the potential interplay between genes and the environment and all of smaller scales. All studies included persons born in Denmark May 1981 or later. Collectively, these forerunners informed on the best possible study design to be used in the iPSYCH2012. In short, the very first selected 413 persons with schizophrenia like psychoses (N=186) and affective disorders (N=256) diagnosed up until October 1999. For each case, the samples in the Danish Neonatal Screening Biobank physically located immediately before and after the sample were selected as controls^1^. The second approach (SKIZO2005) utilized persons diagnosed with schizophrenia and bipolar affective disorder up until September 2005 first including 231 genetic markers and later whole genome genotyped^2–5^. The third approach (SKIZO2006) utilized all persons diagnosed with schizophrenia up until December 31, 2006 reusing biological material already collected in the previous study^5–7^. Finally, SKIZO2008 utilising new persons diagnosed with schizophrenia up until December 31, 2008 ^8^. Controls for SKIZO2005, SKIZO2006 and SKIZO2008 were individually time-matched using the following criteria, born in Denmark, had the same birthday as the corresponding case, and were alive and not diagnosed with schizophrenia when the case was first diagnosed. Genome wide genotyping was performed using the Illumina Human 610-quad beadchip. As demonstrated most persons diagnosed with schizophrenia up until 31 December 2008 had already been genotyped prior to the iPSYCH2012 sample, and was therefore not included here. The latter three designs include a total of 1887 cases of schizophrenia and 1863 individually time-matched controls. These individually matched designs are capable of estimating unbiased incidence rate ratios of the exposures/markers of interest^9^, but challenging in situations where the matching is broken, e.g., due to missing information in the Danish Neonatal Screening Biobank or failure in genotyping/QC. The increased capability for the iPSYCH2012 sample including 30,000 randomly samples cohort members enabled us the unique possibility to introduce a large case cohort sample superior to previous studies while allowing inclusion of previously genotyped samples. Although, inclusion of previously genotyped cases is methodologically straightforward, inclusion of previously genotyped controls is challenging as they are conditioned to be alive and without schizophrenia at different points in time (when then corresponding case was first diagnosed). The iPSYCH2012 case-cohort sample provides the rare opportunity to analyse genetic data and environmental data respecting the longitudinal nature of human life^10^.

## Supplementary text 2

Both the GenCall and Birdseed call sets were filtered to variants that had a call frequency greater than 97% and a Hardy-Weinberg Equilibrium (HWE) p-value greater than 1 × 10^-6^. Only females were used to compute the HWE p-value for X-chromosome variants. The filtered GenCall and Birdseed datasets were merged together by taking a consensus call between the two algorithms. For example, if the two algorithms made different genotype calls or they both had missing calls, the genotype was set to missing. Otherwise, if one algorithm made a call, but the other did not, the non-missing call was used. After merging the GenCall and Birdseed filtered data, any variants that had a call frequency less than 97% or a HWE p-value less than 1 × 10^-6^ were filtered out.

Next, we identified variants with MAF < 0.01 in the merged dataset and that passed the variant filters in the GenCall dataset to create a subset of the zCall dataset for passing rare variants. The zCall subset was filtered such that all variants had a call frequency greater than 97%, HWE p-value < 1 × 10^-6^, and a MAF ≤ 0.05. Lastly, we created a final genotype call set by merging the filtered GenCall/Birdseed call set with the filtered zCall call set.

**Supplementary Table 1.** All probe sequences within the original Illumina manifest were queried against a HG19 database using a nucleotide version of the Basic Local Alignment Search Tool (BLASTN). BLASTN was run with p-value cut-off for matches at 1×10^-15^, first a word length of 30bp was used and probes failing to generate any hits were re-queued without word length limitations. The mappings were compared to three others available in all possible configurations using; 1: **BLASTN:** the procedure described here; 2: **Manifest:** The original positions, “PsychChip_15048346_B.csv”*; 3: **Update:** The Illumina “InfiniumPsychArray-24v1-1_A1.csv_Physical-and-Genetic-Coordinates.txt”╪; 4: **Broad** Institute in house reference.

Each comparison was assigned “pass” if the mappings are concordant, “fail” if discordant or “unknown” if one of the mappings contained no information. Each observed permutation was evaluated and assigned a mapping. The resulting mappings updates are available upon request.

* https://www.med.unc.edu/pgc/files/resultfiles/psychchip_15048346_b.csv.zip

**╪**ftp://webdata:webdata@ussd-ftp.illumina.com/Downloads/ProductFiles/PsychArray/v1-1/infinium-psycharray-24-v1-1-a1-physical-and-genetic-coordinates.zip

| Number | BLASTN:Manifest | BLASTN:Update | BLAST:Broad | Manifest:Update | Manifest:Broad | Update:Broad | % of probes | Mapping used |
| --- | --- | --- | --- | --- | --- | --- | --- | --- |
| 1 | pass | pass | pass | pass | pass | pass | 95.1230% | All agree |
| 2 | pass | unknown | pass | unknown | pass | unknown | 2.2192% | BLASTN |
| 3 | pass | pass | fail | pass | fail | fail | 1.3680% | Manifest |
| 4 | pass | fail | pass | fail | pass | fail | 0.4987% | Manifest |
| 5 | fail | pass | pass | fail | fail | pass | 0.4229% | Broad |
| 6 | pass | unknown | fail | unknown | fail | unknown | 0.1747% | Manifest |
| 7 | fail | pass | fail | fail | fail | fail | 0.0549% | Manifest |
| 8 | fail | unknown | pass | unknown | fail | unknown | 0.0462% | Broad |
| 9 | fail | pass | fail | fail | pass | fail | 0.0280% | Update |
| 10 | fail | fail | pass | pass | fail | fail | 0.0246% | Update |
| 11 | fail | fail | fail | pass | fail | fail | 0.0121% | Manifest |
| 12 | fail | unknown | fail | unknown | fail | unknown | 0.0079% | BLASTN |
| 13 | fail | fail | fail | pass | pass | pass | 0.0069% | Broad |
| 14 | fail | unknown | fail | unknown | pass | unknown | 0.0049% | Broad |
| 15 | pass | fail | fail | fail | fail | pass | 0.0035% | Broad |
| 16 | pass | fail | fail | fail | fail | fail | 0.0025% | Manifest |
| 17 | fail | fail | fail | fail | pass | fail | 0.0010% | Manifest |
| 18 | fail | fail | fail | fail | fail | fail | 0.0008% | Chr 0, pos 0 |
| 19 | fail | fail | fail | fail | fail | pass | 0.0002% | Broad |

## Reference List

1 Sullivan P. F., Daly M. J. & O'Donovan M. Genetic architectures of psychiatric disorders: the emerging picture and its implications. Nature reviews. Genetics 13, 537–551, doi:10.1038/nrg3240 (2012).

2 O'Donovan M. C. & Owen M. J. The implications of the shared genetics of psychiatric disorders. Nat Med 22, 1214–1219, doi:10.1038/nm.4196 (2016).

3 Sullivan P. F. et al. Psychiatric Genomics: An Update and an Agenda. bioRxiv, doi:10.1101/115600 (2017).

4 McGrath J. J., Mortensen P. B., Visscher P. M. & Wray N. R. Where GWAS and epidemiology meet: opportunities for the simultaneous study of genetic and environmental risk factors in schizophrenia. Schizophrenia bulletin 39, 955–959, doi:10.1093/schbul/sbt108 (2013).

5 Frank L. Epidemiology. When an entire country is a cohort. Science 287, 2398–2399 (2000).

6 Schwartz S. & Susser E. The use of well controls: an unhealthy practice in psychiatric research. Psychological medicine 41, 1127–1131, doi:10.1017/S0033291710001595 (2011).

7 Schwartz S. & Susser E. Genome-wide association studies: does only size matter? The American journal of psychiatry 167, 741–744, doi:167/7/741 [pii];10.1176/appi.ajp.2010.10030465 [doi] (2010).

8 Borgan O., Langholz B., Samuelsen S. O., Goldstein L. & Pogoda J. Exposure stratified case-cohort designs. Lifetime Data Anal 6, 39–58 (2000).

9 Clayton D. & Hills M. Statistical models in epidemiology. (Oxford University Press, y1993).

10 Mortensen P. B. et al. Toxoplasma gondii as a risk factor for early-onset schizophrenia: analysis of filter paper blood samples obtained at birth. Biol. Psychiatry 61, 688–693 (2007).

11 Nyegaard M. et al. CACNA1C (rs1006737) is associated with schizophrenia. Molecular psychiatry 15, 119–121, doi:10.1038/mp.2009.69 (2010).

12 Mortensen P. B. et al. Neonatal antibodies to infectious agents and risk of bipolar disorder: a population-based case-control study. Bipolar disorders 13, 624–629, doi:10.1111/j.1399-5618.2011.00962.x (2011).

13 Mortensen P. B. et al. A Danish National Birth Cohort study of maternal HSV-2 antibodies as a risk factor for schizophrenia in their offspring. Schizophrenia research 122, 257–263, doi:10.1016/j.schres.2010.06.010 (2010).

14 Demontis D. et al. Association of GRIN1 and GRIN2A-D with schizophrenia and genetic interaction with maternal herpes simplex virus-2 infection affecting disease risk. American journal of medical genetics. Part B, Neuropsychiatric genetics : the official publication of the International Society of Psychiatric Genetics 156B, 913–922, doi:10.1002/ajmg.b.31234 (2011).

15 Pedersen C. B. et al. Risk of schizophrenia in relation to parental origin and genome-wide divergence. Psychological medicine 42, 1515–1521, doi:10.1017/S0033291711002376 (2012).

16 Mortensen P. B. et al. Maternal Antibodies to Cytomegalovirus and Schizophrenia Risk. Schizophrenia bulletin 37, 58–58 (2011).

17 Borglum A. D. et al. Genome-wide study of association and interaction with maternal cytomegalovirus infection suggests new schizophrenia loci. Molecular psychiatry 19, 325–333, doi:10.1038/mp.2013.2 (2014).

18 Pedersen C. B., Gotzsche H., Moller J. O. & Mortensen P. B. The Danish Civil Registration System. A cohort of eight million persons. Danish medical bulletin 53, 441–449 (2006).

19 Mors O., Perto G. P. & Mortensen P. B. The Danish Psychiatric Central Research Register. Scand. J. Public Health 39, 54–57 (2011).

20 Organization W. H. WHO ICD-10: Psykiske lidelser og adfærdsmæssige forstyrrelser. Klassifikation og diagnosekriterier [WHO ICD-10: Mental and Behavioural Disorders. Classification and Diagnostic Criteria]. (Munksgaard Danmark, 1994).

21 Waltoft B. L., Pedersen C. B., Nyegaard M. & Hobolth A. The importance of distinguishing between the odds ratio and the incidence rate ratio in GWAS. BMC Med Genet 16, 71, doi:10.1186/s12881-015-0210-1 (2015).

22 Norgaard-Pedersen B. & Hougaard, D. M. Storage policies and use of the Danish Newborn Screening Biobank. J. Inherit. Metab Dis 30, 530–536 (2007).

23 Poulsen J. B. et al. High-Quality Exome Sequencing of Whole-Genome Amplified Neonatal Dried Blood Spot DNA. PloS one 11, e0153253, doi:10.1371/journal.pone.0153253 (2016).

24 Hollegaard M. V., Grauholm J., Norgaard-Pedersen B. & Hougaard D. M. DNA methylome profiling using neonatal dried blood spot samples: a proof-of-principle study. Mol Genet Metab 108, 225–231, doi:10.1016/j.ymgme.2013.01.016 (2013).

25 Eyles D. W. et al. The utility of neonatal dried blood spots for the assessment of neonatal vitamin D status. Paediatric and perinatal epidemiology 24, 303–308, doi:10.1111/j.1365-3016.2010.01105.x (2010).

26 Skogstrand K. et al. Simultaneous measurement of 25 inflammatory markers and neurotrophins in neonatal dried blood spots by immunoassay with xMAP technology. Clinical chemistry 51, 1854-1866, doi:10.1373/clinchem.2005.052241 (2005).

27 Bybjerg-Grauholm J. et al. RNA sequencing of archived neonatal dried blood spots. Molecular genetics and metabolism reports 10, 33–37, doi:10.1016/j.ymgmr.2016.12.004 (2017).

28 Hollegaard M. V. et al. Genome-wide scans using archived neonatal dried blood spot samples. BMC. Genomics 10, 297 (2009).

29 Hollegaard M. V. et al. Robustness of genome-wide scanning using archived dried blood spot samples as a DNA source. BMC genetics 12, 58, doi:10.1186/1471-2156-12-58 (2011).

30 Gunderson K. L. et al. Whole-Genome Genotyping. 410, 359–376, doi:10.1016/s0076-6879(06)10017-8 (2006).

31 Illumina. Illumina GenCall Data Analysis Software. Illumina Techinal Note, <https://www.illumina.com/Documents/products/technotes/technotegencalldataanalysissoftware.pdf> (2005).

32 Korn J. M. et al. Integrated genotype calling and association analysis of SNPs, common copy number polymorphisms and rare CNVs. Nature genetics 40, 1253–1260, doi:10.1038/ng.237 (2008).

33 Goldstein J. I. et al. zCall: a rare variant caller for array-based genotyping: genetics and population analysis. Bioinformatics 28, 2543–2545, doi:10.1093/bioinformatics/bts479 (2012).

34 Chang C. C. et al. Second-generation PLINK: rising to the challenge of larger and richer datasets. GigaScience 4, 7, doi:10.1186/s13742-015-0047-8 (2015).

35 Purcell S. et al. PLINK: A tool set for whole-genome association and population-based linkage analyses. Am J Hum Genet 81, 559–575 (2007).

36 Hartlev M. Genomic Databases and Biobanks in Denmark. J Law Med Ethics 43, 743–753, doi:10.1111/jlme.12316 (2015).

37 Statistics D. Guidelines for Transferring Aggregated Results from Statistics Denmark's Research Services, <http://dst.dk/ext/8804839804/0/forskning/Guidelines-for-Transferring-Aggregated-Results-from-Statistics-Denmark-s-Research-Services--pdf> (2017).

38 Price A. L. et al. Long-range LD can confound genome scans in admixed populations. Am J Hum Genet 83, 132–135; author reply 135-139, doi:10.1016/j.ajhg.2008.06.005 (2008).

39 Gogarten S. M. et al. GWASTools: an R/Bioconductor package for quality control and analysis of genome-wide association studies. Bioinformatics 28, 3329–3331, doi:10.1093/bioinformatics/bts610 (2012).

40 Patterson N., Price A. L. & Reich D. Population structure and eigenanalysis. Plos Genet 2, e190, doi:10.1371/journal.pgen.0020190 (2006).

41 Cantor-Graae E. & Pedersen C. B. Risk of schizophrenia in second-generation immigrants: a Danish population-based cohort study. Psychological medicine 37, 485–494, doi:10.1017/S0033291706009652 (2007).

42 Pedersen C. B. No evidence of time trends in the urban-rural differences in schizophrenia risk among five million people born in Denmark from 1910 to 1986. Psychological medicine 36, 211–219, doi:10.1017/S003329170500663X (2006).

43 McGrath J. J. et al. A comprehensive assessment of parental age and psychiatric disorders. JAMA psychiatry 71, 301–309, doi:10.1001/jamapsychiatry.2013.4081 (2014).

44 Dean K. et al. Full spectrum of psychiatric outcomes among offspring with parental history of mental disorder. Archives of general psychiatry 67, 822–829, doi:10.1001/archgenpsychiatry.2010.86 (2010).

45 Pedersen C. B. et al. A comprehensive nationwide study of the incidence rate and lifetime risk for treated mental disorders. JAMA psychiatry 71, 573–581, doi:10.1001/jamapsychiatry.2014.16 (2014).

46 McGrath J. J. et al. Neonatal vitamin D status and risk of schizophrenia: a population-based case-control study. Archives of general psychiatry 67, 889–894, doi:10.1001/archgenpsychiatry.2010.110 (2010).

47 Ostergaard S. D. et al. Predicting ADHD by Assessment of Rutter's Indicators of Adversity in Infancy. PloS one 11, doi:ARTN e0157352 10.1371/journal.pone.0157352 (2016).

48 Wimberley T. et al. Predictors of treatment resistance in patients with schizophrenia: a population-based cohort study. Lancet Psychiatry 3, 358–366, doi:10.1016/S2215-0366(15)00575-1 (2016).

49 Dalsgaard S., Leckman J. F., Mortensen P. B., Nielsen H. S. & Simonsen M. Effect of drugs on the risk of injuries in children with attention deficit hyperactivity disorder: a prospective cohort study. Lancet Psychiatry 2, 702–709, doi:10.1016/S2215-0366(15)00271-0 (2015).

50 Nordentoft M., Mortensen P. B. & Pedersen C. B. Absolute risk of suicide after first hospital contact in mental disorder. Archives of general psychiatry 68, 1058–1064, doi:10.1001/archgenpsychiatry.2011.113 (2011).

51 Dalsgaard S., Ostergaard S. D., Leckman J. F., Mortensen P. B. & Pedersen M. G. Mortality in children, adolescents, and adults with attention deficit hyperactivity disorder: a nationwide cohort study. Lancet 385, 2190–2196, doi:10.1016/S0140-6736(14)61684-6 (2015).

52 Gjerstorff M. L. The Danish Cancer Registry. Scand J Public Health 39, 42–45, doi:10.1177/1403494810393562 (2011).

53 Kildemoes H. W., Sorensen H. T. & Hallas J. The Danish National Prescription Registry. Scand J Public Health 39, 38–41, doi:10.1177/1403494810394717 (2011).

54 Lynge E., Sandegaard J. L. & Rebolj M. The Danish National Patient Register. Scand. J. Public Health 39, 30–33, doi:39/7_suppl/30 [pii];10.1177/1403494811401482 [doi] (2011).

55 Baadsgaard M. & Quitzau J. Danish registers on personal income and transfer payments. Scand J Public Health 39, 103–105, doi:10.1177/1403494811405098 (2011).

56 Jensen V. M. & Rasmussen A. W. Danish Education Registers. Scand J Public Health 39, 91–94, doi:10.1177/1403494810394715 (2011).

57 Petersson F., Baadsgaard M. & Thygesen L. C. Danish registers on personal labour market affiliation. Scand J Public Health 39, 95–98, doi:10.1177/1403494811408483 (2011).

58 Schizophrenia Working Group of the Psychiatric Genomics, C. Biological insights from 108 schizophrenia-associated genetic loci. Nature 511, 421–427, doi:10.1038/nature13595 (2014).

59 Meier S. M. et al. High loading of polygenic risk in cases with chronic schizophrenia. Molecular psychiatry, doi:10.1038/mp.2015.130 (2015).

60 Wray N. R., Goddard M. E. & Visscher P. M. Prediction of individual genetic risk to disease from genome-wide association studies. Genome Res 17, 1520–1528, doi:10.1101/gr.6665407 (2007).

61 Purcell S. M. et al. Common polygenic variation contributes to risk of schizophrenia and bipolar disorder. Nature 460, 748–752 (2009).

62 Dudbridge F. Polygenic Epidemiology. Genet Epidemiol 40, 268–272, doi:10.1002/gepi.21966 (2016).

63 Agerbo E. et al. Modelling the contribution of family history and variation in single nucleotide polymorphisms to risk of schizophrenia: a Danish national birth cohort-based study. Schizophrenia research 134, 246–252, doi:10.1016/j.schres.2011.10.025 (2012).

64 Agerbo E. et al. Polygenic Risk Score, Parental Socioeconomic Status, Family History of Psychiatric Disorders, and the Risk for Schizophrenia: A Danish Population-Based Study and Meta-analysis. JAMA psychiatry 72, 635–641, doi:10.1001/jamapsychiatry.2015.0346 (2015).

65 Benros M. E. et al. Influence of Polygenic Risk Scores on the Association Between Infections and Schizophrenia. Biol Psychiatry 80, 609–616, doi:10.1016/j.biopsych.2016.04.008 (2016).

66 Wimberley T. et al. Polygenic Risk Score for Schizophrenia and Treatment-Resistant Schizophrenia. Schizophr Bull, doi:10.1093/schbul/sbx007 (2017).

67 Laursen T. M. et al. Association of the polygenic risk score for schizophrenia with mortality and suicidal behavior - A Danish population-based study. Schizophrenia research, doi:10.1016/j.schres.2016.12.001 (2016).

68 Demyttenaere K. et al. Prevalence, severity, and unmet need for treatment of mental disorders in the World Health Organization World Mental Health Surveys. Jama 291, 2581–2590, doi:10.1001/jama.291.21.2581 [doi];291/21/2581 [pii] (2004).

69 Bock C., Bukh J. D., Vinberg M., Gether U. & Kessing L. V. Validity of the diagnosis of a single depressive episode in a case register. Clin. Pract. Epidemiol. Ment. Health 5, 4 (2009).

70 Kessing L. V. Validity of diagnoses and other clinical register data in patients with affective disorder. European Psychiatry 13, 392–398 (1998).

71 Lauritsen M. B. et al. Validity of childhood autism in the Danish Psychiatric Central Register: findings from a cohort sample born 1990-1999. J. Autism Dev. Disord 40, 139–148 (2010).

72 Uggerby P., Ostergaard S. D., Roge R., Correll C. U. & Nielsen J. The validity of the schizophrenia diagnosis in the Danish Psychiatric Central Research Register is good. Dan. Med J 60, A4578, doi:A4578 [pii] (2013).

73 Jakobsen K. D. et al. Reliability of clinical ICD-10 schizophrenia diagnoses. Nord. J. Psychiatry 59, 209–212 (2005).

74 Mohr-Jensen C., Vinkel Koch S., Briciet Lauritsen M. & Steinhausen H. C. The validity and reliability of the diagnosis of hyperkinetic disorders in the Danish Psychiatric Central Research Registry. Eur Psychiatry 35, 16–24, doi:10.1016/j.eurpsy.2016.01.2427 (2016).

75 Frank L. Epidemiology. The epidemiologist's dream: Denmark. Science 301, 163, doi:10.1126/science.301.5630.163 (2003).

## Reference List

1 Mortensen P. B. et al. Toxoplasma gondii as a risk factor for early-onset schizophrenia: analysis of filter paper blood samples obtained at birth. Biol. Psychiatry 61, 688–693 (2007).

2 Nyegaard M. et al. CACNA1C (rs1006737) is associated with schizophrenia. Molecular psychiatry 15, 119–121, doi:10.1038/mp.2009.69 (2010).

3 Mortensen P. B. et al. Neonatal antibodies to infectious agents and risk of bipolar disorder: a population-based case-control study. Bipolar disorders 13, 624–629, doi:10.1111/j.1399-5618.2011.00962.x (2011).

4 Mortensen P. B. et al. A Danish National Birth Cohort study of maternal HSV-2 antibodies as a risk factor for schizophrenia in their offspring. Schizophrenia research 122, 257–263, doi:10.1016/j.schres.2010.06.010 (2010).

5 Demontis D. et al. Association of GRIN1 and GRIN2A-D with schizophrenia and genetic interaction with maternal herpes simplex virus-2 infection affecting disease risk. American journal of medical genetics. Part B, Neuropsychiatric genetics : the official publication of the International Society of Psychiatric Genetics 156B, 913–922, doi:10.1002/ajmg.b.31234 (2011).

6 Pedersen C. B. et al. Risk of schizophrenia in relation to parental origin and genome-wide divergence. Psychological medicine 42, 1515–1521, doi:10.1017/S0033291711002376 (2012).

7 Mortensen P. B. et al. Maternal Antibodies to Cytomegalovirus and Schizophrenia Risk. Schizophrenia bulletin 37, 58–58 (2011).

8 Borglum A. D. et al. Genome-wide study of association and interaction with maternal cytomegalovirus infection suggests new schizophrenia loci. Molecular psychiatry 19, 325–333, doi:10.1038/mp.2013.2 (2014).

9 Breslow N. E. & Day N. E. Statistical methods in cancer research. 1. The analysis of case-control studies. (International Agency for Research on Cancer, 1980).

10 Waltoft B. L., Pedersen C. B., Nyegaard M. & Hobolth A. The importance of distinguishing between the odds ratio and the incidence rate ratio in GWAS. BMC Med Genet 16, 71, doi:10.1186/s12881-015-0210-1 (2015).

